# Diminished adherence of *Biomphalaria glabrata* embryonic cell line to sporocysts of *Schistosoma mansoni* following programmed knockout of the allograft inflammatory factor

**DOI:** 10.1101/2020.04.07.029629

**Authors:** Fernanda Sales Coelho, Rutchanee Rodpai, André Miller, Shannon E. Karinshak, Victoria H. Mann, Omar dos Santos Carvalho, Roberta Lima Caldeira, Marina de Moraes Mourão, Paul J. Brindley, Wannaporn Ittiprasert

## Abstract

**Background:** Larval development in an intermediate host gastropod snail of the genus *Biomphalaria* is an obligatory component of the life cycle of *Schistosoma mansoni*. Understanding of the mechanism(s) of host defense may hasten the development of tools that block transmission of schistosomiasis. The allograft inflammatory factor 1, AIF, which is evolutionarily conserved and expressed in phagocytes, is a marker of macrophage activation in both mammals and invertebrates. AIF enhances cell proliferation and migration. The embryonic cell line, termed Bge, from *Biomphalaria glabrata* is a versatile resource for investigation of the snail-schistosome relationship since Bge exhibits a hemocyte-like phenotype. Hemocytes perform central roles in innate and cellular immunity in gastropods and in some cases can kill the parasite. However, the Bge cells do not kill the parasite *in vitro*.

**Methods:** Bge cells were transfected by electroporation with plasmid pCas*-Bg*AIFx4, encoding the Cas9 nuclease and a guide RNA specific for exon 4 of the *B. glabrata* AIF (*Bg*AIF) gene. Transcript levels for Cas9 and for *Bg*AIF were monitored by quantitative reverse-transcription-PCR and, in parallel, adhesion of gene-edited Bge cells during co-culture with of schistosome sporocysts was assessed.

**Results:** Gene knockout manipulation induced gene-disrupting indels, frequently 1-2 bp insertions and/or 8-30 bp deletions, at the programmed target site; a range from 9 to 17% of the copies of the *Bg*AIF gene in the Bge population of cells were mutated. Transcript levels for *Bg*AIF were reduced by up to 73% (49.5±20.2% S.D, *P* ≤ 0.05, n =12). Adherence by *Bg*AIF gene-edited (Δ*Bg*AIF) Bge to sporocysts diminished in comparison to wild type cells, although cell morphology did not change. Specifically, as scored by a semi-quantitative cell adherence index (CAI), fewer Δ*Bg*AIF than control wild type cells adhered to sporocysts; control CAI, 2.66±0.10, Δ*Bg*AIF, 2.30±0.22 (*P* ≤ 0.01).

**Conclusion:** The findings supported the hypothesis that *Bg*AIF plays a role in the adherence of *B. glabrata* hemocytes to sporocysts during schistosome infection *in vitro*. This demonstration of the activity of programmed gene editing will enable functional genomics approaches using CRISPR/Cas9 to investigate additional components of the snail-schistosome host-parasite relationship.

## Background

Evolution endowed the schistosomes with a complex life cycle that includes both a freshwater gastropod intermediate host and a definitive mammalian host. Several species of the freshwater snail genus *Biomphalaria* are the intermediate host for *Schistosoma mansoni*. The neotropical species, *Biomphalaria glabrata* has been studied extensively with respect to host-parasite relationship and coevolution with *S. mansoni* especially on mechanisms of susceptibility and/or resistance to the compatible parasites (1, 2). Genetic variation is evident among isolates and strains of *B. glabrata*, both in the laboratory and in the field, resulting in a spectrum of the susceptibility of infection with *S. mansoni* (3). Considerable advances have been made in the exploration and characterization of mechanisms of the internal defenses system (IDS) of the snail that determine susceptibility and resistance to schistosome (4-11). The resistance phenotype is underpinned by a complex genetic trait, where the schistosome larva fails to develop as the consequence of innate and cellular immune responses. Hemocytes of resistant snails encapsulate and destroy the sporocyst (11-18).

*B. glabrata* embryonic cell line (Bge) (19) remains to date the only established cell line from any mollusk. The cell line originates from five-day-old embryos of *B. glabrata* susceptible to infection with *S. mansoni*. The Bge cell line has been studied extensively to interrogate the host-parasite relationship because the Bge cell exhibits a hemocyte-like behavior that includes encapsulation of the larval parasite, but does not kill the parasites (20-28).

The genome sequence of *B. glabrata* has been reported (29), along with ongoing transcriptome and proteome catalogues that include factors participating in immunological surveillance, phagocytosis, cytokine responses, and pathogen recognition receptor elements including Toll-like receptors and fibrinogen-related proteins (30-36). An orthologue of the evolutionary conserved allograft inflammatory factor (AIF) is an evolutionary conserved protein typically expressed in phagocytes and granular leukocytes in both vertebrate and invertebrate.

Functions demonstrated for AIF include macrophage activation, enhancement of cellular proliferation and of migration in mammalian and invertebrate cells; protostomes and deuterostomes (37-41). AIF also plays a key role in the protective response by *B. glabrata* to invasion by schistosomes (8, 9). *Bg*AIF, the orthologue in *B. glabrata* is expressed in hemocytes, which participate in phagocytosis, cellular proliferation, and cellular migration. Elevated expression of *Bg*AIF is a characteristic of the resistance of *B. glabrata* to schistosome infection and has been considered as a marker of hemocyte activation (8, 9).

Expression of AIF also is seen during hemocyte activation in oysters (36, 38, 42, 43) and during hepatic inflammation during murine schistosomiasis (44, 45). We hypothesized that *Bg*AIF was involved in cell mediated immune response(s) by *B. glabrata* through activation of hemocyte cell adhesion and/or migration after the schistosome miracidium has penetrated into the tissues of the snail. We addressed this hypothesis by using CRISPR/Cas9-based programmed genome editing to interrupt the *Bg*AIF gene of *B. glabrata* in the Bge cell line, following reports that indicated the utility of using CRISPR-based programmed gene knockout approach in other mollusks including the Pacific oyster, *Crassostrea gigas* and the slipper limpet, *Crepidula fornicata* and the gastropod, *Lymnaea stagnalis* (46-48). As detailed below, we demonstrated the activity of programmed genome editing in Bge cells, with gene knockout at the *Bg*AIF locus.

## Methods

### Gene editing construct

The gene encoding the allograft inflammatory factor of *B. glabrata, Bg*AIF (2,226 bp; accession number BGLB005061, https://www.vectorbase.org/) includes five exons interrupted by four introns (Fig. 1a). A guide RNA (gRNA) for Cas9-catalyzed gene editing specific for the target *B. glabrata* gene locus, *Bg*AIF, was identified in the BGLB005061 sequence using the ‘CHOPCHOP’ v3 tool, https://chopchop.cbu.uib.no/, with default parameters compatible for the protospacer adjacent motif, NGG, of Cas9 from *Streptococcus pyogenes* (49-51) and screened for off-target sites against the *Biomphalaria glabrata* genome (29). Based on the guidance from the CHOPCHOP analysis, we chose the top ranked guide RNA (gRNA), AGACTTTGTTAGGATGATGC, specific for exon 4 of the AIF gene, with predicted high CRISPR/Cas9 efficiency for double-stranded cleavage in tandem with an absence of off-target activity in the genome of *B. glabrata* (Fig. 1a). A CRISPR/Cas9 vector encoding the gRNA targeting exon 4 of *Bg*AIF under the control of the mammalian U6 promoter and encoding Cas 9, with nuclear localization signals 1 and 2, driven by the human cytomegalovirus (CMV) immediate early enhancer and promoter was assembled using the GeneArt CRISPR Nuclease Vector system (Thermo Fisher Scientific, Waltham, MA), according to the manufacturer’s protocol. Briefly, the 20 nt of either target (including ‘gtttt’ on the 3’ end) or complementary to target (including ‘cggtg’ on the 3’end) sequences were synthesized commercially, and annealed according to manufacturer’s protocol. The annealed double strand DNA (dsDNA) was ligated into the linearized GeneArt^®^ CRISPR Nuclease vector via *BamH*I and *BsmB*I restriction sites, respectively, and the construct was termed pCas-*Bg*AIFx4 (Fig. 1b). (The sequence of GeneArt CRISPR nuclease vector backbone is available at https://www.thermofisher.com/order/catalog/product/A21174#/A21174). Chemically competent TOP10, *E. coli* cells (Invitrogen, Thermo Fisher Scientific) were transformed with pCas-*Bg*AIFx4 by the heat shock method and cultured on LB-agar supplemented with ampicillin at 100 µg/ml. Subsequently, the integrity of the recombinant plasmids from several single colonies of ampicillin-resistant *E. coli* transformants was confirmed by amplicon PCR-based Sanger direct nucleotide sequence analysis using a U6 gene-specific primer for gRNA ligation and orientation (Fig. 1b).

**Fig. 1.**
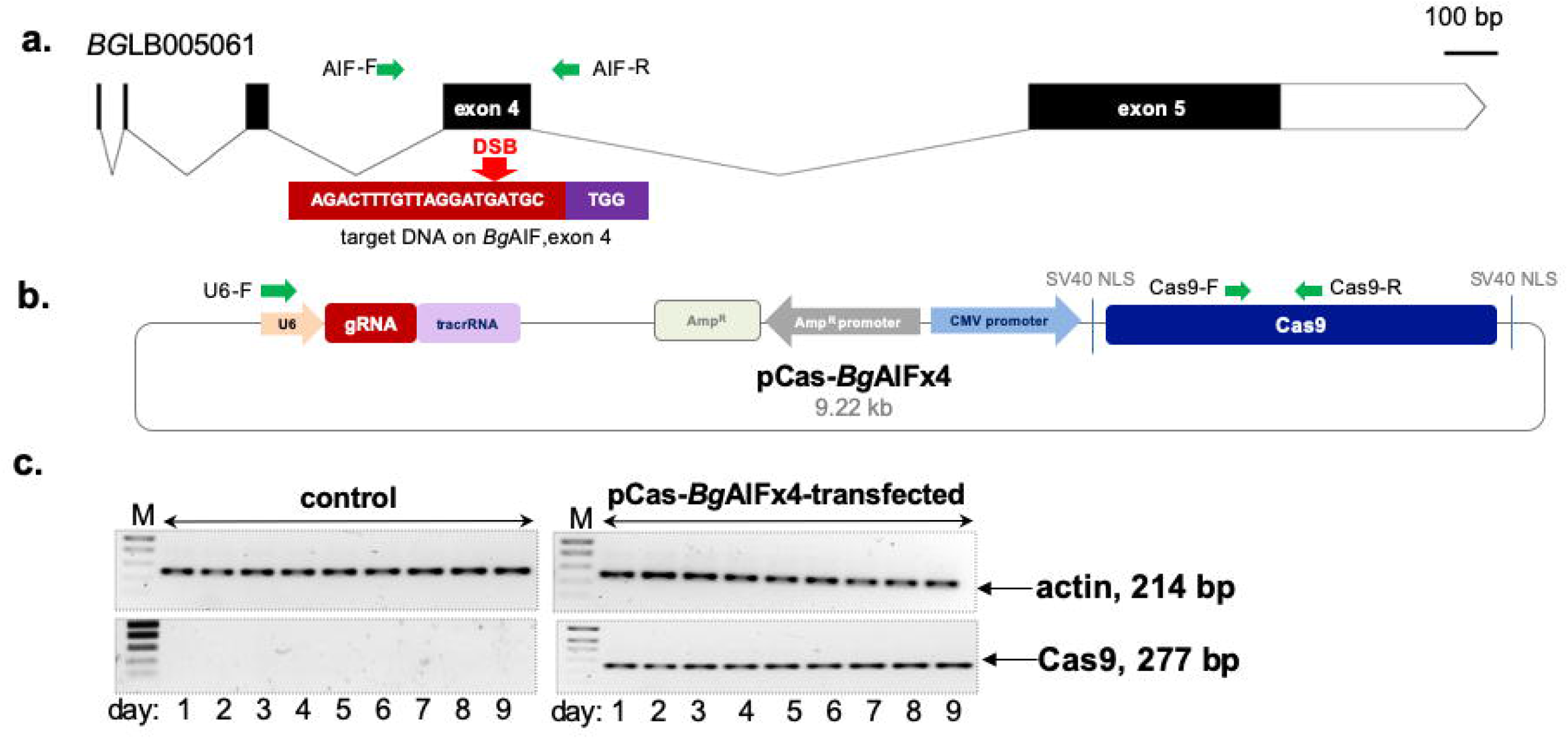
Schematic diagram of *Bg*AIF gene structure, CRISPR/Cas9 vector and expression in Bge cell. **a** Gene structure of *B. glabrata* allograft inflammatory factor (*Bg*AIF), accession number BGLB005061 and gene editing target locus (red box) on exon 4. *Bg*AIF gene composed of 5 exons and 4 introns. The green arrows indicate the location of primers flanking expected double strand breaks (DSB) which were used in PCR to generate the on-target amplicon for INDELs estimation. **b** Map of the pCas-*Bg*AIFx4 vector which includes the Pol III-dependent mammalian U6 gene promoter (red arrow) to drive transcription of the guide RNA targeting exon 4 of *Bg*AIF gene (red arrow) and the CMV promoter to drive expression of the *S. pyogenes* Cas9 nuclease (blue arrow). Primer pairs specific for the guide RNA and for Cas9 are indicated (green arrows). **c** Expression of Cas9 and of *Bg*Actin (as the reference gene) transcripts as established by semi-quantitative RT-PCR in pCas-*Bg*AIF-transfected (right) and control (left) Bge cells from days one to nine following transfection. The amplicons of the expected sizes are as indicated: 23 bp for Cas9 and 214 bp for *Bg*Actin. All RNA samples were positive for the *Bg*Actin reference gene; the 214 bp band.

### *Biomphalaria glabrata* embryonic (Bge) cell line culture

The Bge cell line was provided by the Schistosomiasis Resource Center (SRC), Biomedical Research Institute (BRI), Rockville, MD. Historically, the Bge cell line was sourced by the SRC from the American Type Culture Collection (Manassas, VA), catalog no. ATCC CRL 1494, and thereafter maintained at BRI for >10 years. Bge cells were maintained at 26°C in air in ‘Bge medium’, which is comprised of 22% (v/v) Schneider’s *Drosophila* medium, 0.13% galactose, 0.45% lactalbumin hydrolysate, 0.5% (v/v) phenol red solution, 20 µg/ml gentamycin, and supplemented with 10% heat-inactivated fetal bovine serum (24, 52). Bge cells were grown to 80% confluence before transfection by electroporation with pCas-*Bg*AIFx4. The Bge cells were free of contamination with *Mycoplasma*, as established with a PCR-based test (LookOut® Mycoplasma PCR Detection kit, Sigma-Aldrich, St. Louis, MO).

### Transfection of Bge cells by square wave electroporation

Bge cells were harvested using a cell scraper, washed twice in Bge medium, counted, and resuspended at 20,000 cell/µl in Opti-MEM medium (Sigma-Aldrich, St. Louis, MO). Two million cells were transferred into 0.2 mm path length electroporation cuvettes (BTX Harvard Apparatus, Hollister, MA) containing 6 µg pCas-*Bg*AIFx4 in ∼100 µl Opti-MEM. The cells were subjected to electroporation using one pulse at 125 volts for 20 milliseconds, using a square wave pulse generator (ECM 830, BTX Harvard Apparatus). Immediately thereafter, the Bge cells were maintained in 12-well plates (Greiner Bio-One) at 26°C. The mock control included Opti-MEM only for electroporation. The presence of transcripts encoding the *B. glabrata* actin and the Cas9 was monitored daily for nine days following transfection by electroporation (Fig. 1c).

### Sequential isolation of total RNA and genomic DNA

To monitor the transfection of Bge cell by pCas9-*Bg*AIFx4, we investigated the expression of Cas9 in Bge cells by reverse transcription PCR (RT-PCR). Both total RNA and genomic DNA were extracted sequentially from cell pellets, as described (53, 54). In brief, each sample of total RNA sample was extracted using the RNAzol^®^ RT reagent (Molecular Research Center, Inc., Cincinnati, OH) according to the manufacturer’s protocol. Subsequently, the DNA/protein pellet retained after recovery of RNA was resuspend in DNAzol^®^ solution (Molecular Research Center, Inc), from which total DNA was recovered. The RNAs and DNAs were dissolved in nuclease-free water and their concentration and purity established by spectrophotometry (Nanodrop 1000, Thermo Fisher Scientific).

### Expression of Cas9 in Bge cells

To investigate transcription from the pCas-*Bg*AIFx4 vector following transfection of Bge cells, levels of transcribed Cas9 were investigated by semi-quantitative RT-PCR. The Cas9-specific primers were Cas9-F, 5’-agcatcggccttgatatcgg-3’ and Cas9-R, 5’-agaagctgtcgtccaccttg-3’ (Fig. 1b). Total RNA from the non-transfected, mock (Opti-MEM electroporated-), and pCas-*Bg*AIFx4 DNA electroporated-Bge cells were treated with DNase I (Ambion, Thermo Fisher Scientific) to digest any residual vector pCas-*Bg*AIFx4 DNA and contaminating genomic DNAs. The RNAs were reverse transcribed to cDNA using ProtoScript II reverse transcriptase with oligo dT and random primers (First Strand cDNA Synthesis kit New England Biolabs, Ipswich, MA). RT-PCRs specific for the Cas9 or actin gene of *B. glabrata, Bg*Actin (GenBank accession number U53348.1) were undertaken, with BgActin serving as the positive control for RNA integrity. The primer pairs used for the *Bg*Actin coding sequences were termed actin-F, 5’-aagcgacgttttcttggtgc-3’ and actin-R, 5’-acccataccaaccatcacacc-3’. Amplicons and molecular size standards were separated by electrophoresis through Tris-acetate-EDTA-buffered agarose 1% stained with ethidium bromide (Fig. 1c).

### Analysis of programmed mutation of the allograft inflammatory factor gene of *B. glabrata*

Genomic DNA samples from the mock-transfected and pCas-*Bg*AIFx4-transfected cells were amplified by PCR using AIF-F (5’-gcagatttgcaattcaacactta-3’) and AIF-R (5’-tgccagctagcttactgcat-3’) primers that flank the CRISPR/Cas9 programmed double-stranded break (DSB) site (Fig. 1a). Amplicons of 568 nt in length (from residues 489 to 1,056 of the *Bg*AIF_BGLB0055061 gene) were obtained using the AIF-F and -R primer pair. Amplicons were isolated from the agarose gel using the PCR cleanup and gel extraction kit (ClonTech, Takara USA, Mountain View, CA) and the nucleotide sequence of amplicons determined by Sanger direct sequencing (GENEWIZ, South Plainfield, NJ). Chromatograms of the sequence reads from the control and experimental groups in each replicate experiment were subjected to online analysis using the TIDE algorithm, https://tide.deskgen.com/ (55, 56) and also using the Inference of CRISPR v2 Edits analysis (ICE) software, https://ice.synthego.com/#/ (Synthego Corporation, Redwood City, CA) (57). Estimates of CRISPR efficiency, insertion-deletion (INDEL)-substitution percentages, and the nucleotide sequence of mutant alleles were obtained using both the TIDE and the ICE platforms (55, 56) (Fig. 2a, 2b).

**Fig. 2.**
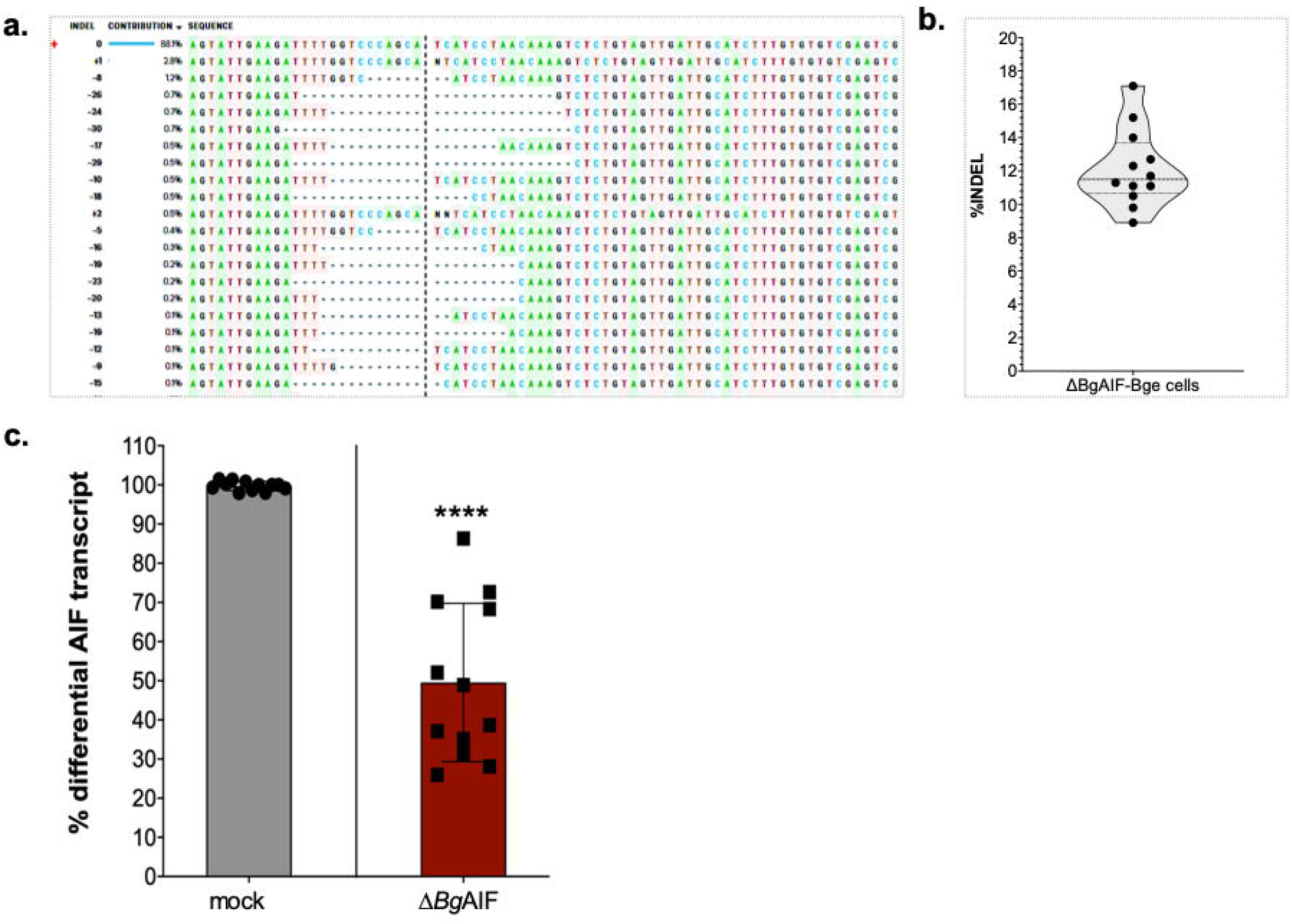
Establishment of *Bg*AIF-knockout lines of Bge cells. **a** Representative examples of frequent gene insertions-deletions (1-2 bp insertions and 8-30 bp deletions, straddling the programmed CRISPR/Cas9-induced double-stranded break in exon 4, as determined by ICE software-based analysis. **b** TIDE algorithm-based violin plot of insertion-deletion percentages (%INDEL) computed using the amplicon sequence traces from the 12 biological replicates of pCas-*Bg*AIF-transfected Bge cell populations. **c** Reduction of *Bg*AIF transcription by about 50% following programmed genome editing of Bge cells (Δ*Bg*AIF-Bge) in comparison to control Bge cells. Mean transcript reduction, 49.55± 20.22 (S.D.) percent, *P* ≤ 0.0001 (****), n =12 (unpaired Student’s *t*-test).

### Quantitative real time PCR analysis of transcription of *Bg*AIF

To evaluate the differential levels of the *Bg*AIF transcript among the control and experimental groups, total RNAs were extracted and treated with DNase I, as above. DNase I treated-RNA (200 ng) was reverse transcribed to cDNA, followed by quantitative RT-PCR, using the ViiA7 Real Time PCR System (Applied Biosystems, Scientific), and the SSoAdvanced Universal SYBR Green Supermix reagents (Bio-Rad), according to the manufacturer’s recommendations. The following nucleotide primers used *Bg*AIF gene-specific primers amplify 119-257 nt of *Bg*AIF GenBank accession number EX001601.1; *Bg*AIF-rt-F, 5’-cctgcttttaacccgacaga-3’ and *Bg*AIF-rt-R, 5’-tgaatgaaagctcctcgtca-3’. Differential *Bg*AIF gene expression were calculated after normalizing with *Bg*Actin (primers as above) and comparison with the non-treated (control) cells. The ΔΔCt method was used to calculate the differential gene expression (58), with assistance of the GraphPad Prism 8 software (San Diego, CA) (Fig. 2c).

### Schistosome sporocysts

Miracidia of the NMRI strain of *S. mansoni* were hatched from eggs recovered from livers of schistosome infected mice (Schistosomiasis Resource Center, Biomedical Research Institute, Rockville, MD) under axenic conditions (28), primary sporocysts were transformed from the miracidia *in vitro*, as described (26). Briefly, miracidia were immobilized by chilling on ice for 25 min, following by pelleting using centrifugation, 500×*g* at 4°C, 60 sec. The miracidia were washed with ice cold Chernin’s balanced salt solution (28 mM NaCl, 0.5mM Na_2_HPO_4_, 2mM KCl, 1.8mM MgSO_4_.7H_2_O, 0.6mM NaHCO_3_, 3.6 mM CaCl_2_.2H_2_O) with 1 mg/ml glucose, trehalose, and antibiotic, 10 µl/ml of 100·× penicillin/streptomycin (Thermo Fisher Scientific), termed CBSS^+^. Approximately 5,000 miracidia per well of 24-well plates were cultured in CBSS+ at 26°C for 24 hrs, after which the sporocysts were washed to remove shed ciliated epidermal plates and other debris, followed by transfer to a 1.5 ml microcentrifuge tube (26).

### Sporocyst-Bge cell binding assay and cell adhesion index (CAI)

To investigate the if *Bg*AIF would affect the ability of cell adhesion to *S. mansoni* sporocyst, we co-cultured the non-transfected Bge cell or non-selected-, transfected-pCas-*Bg*AIFx4 cells (*Bg*AIF depleted-cells named ‘Δ*Bg*AIF-Bge’) with *in vitro* transformed sporocysts, then the cell adhesion index (CAI) were calculated as described (26, 59). With the limitations in this study, we were not be able to select or enrich for *Bg*AIF edited-cells, and hence the Δ*Bg*AIF-Bge cell populations can be considered to be a population of gene mutated mixed with non-modified (wild type) cells. CAI is a semi-quantitative method of cell adhesion to primary sporocysts using the four categories of scores ranging from one to four - lower to higher numbers of cells adherent to the parasite’s surface. In brief, we mixed single cell suspensions of 500,000 Bge cells with 200 freshly prepared-sporocysts (total volume 200 µl of CBSS^+^) in sterile, siliconized tubes (Bio Plas, Thomas Scientific, Swedesboro, NJ). The Bge cell-sporocyst co-culture was maintained at 26°C for 24 hrs. Cellular morphology and adhesion of the cells to the surface of the sporocysts was monitored and recorded using an inverted microscope, at 20× magnification (Zeiss Axio Observer A1, Carl Zeiss LLC, White Plains, NY) after gently transferring the parasite-cell suspension to the tissue culture plate (Greiner Bio-One). Scoring of the adherence index was carried out in a blinded fashion to the investigator reading the score; ≥50 sporocysts from each experimental group were counted each time, and triplicates of each treatment group were scored. Seven independent biological replicates of this CAI-based sporocyst-Bge cell binding assay were carried out. In total, ≥400 sporocysts were examined from each treatment and control group. Averages for the CAI values were calculated from the cell adhesion scores ranging from 1 to 4 (examples presented in Fig. 3a) according to the formula, CAI = total binding value per number of sporocysts (26).

**Fig. 3.**
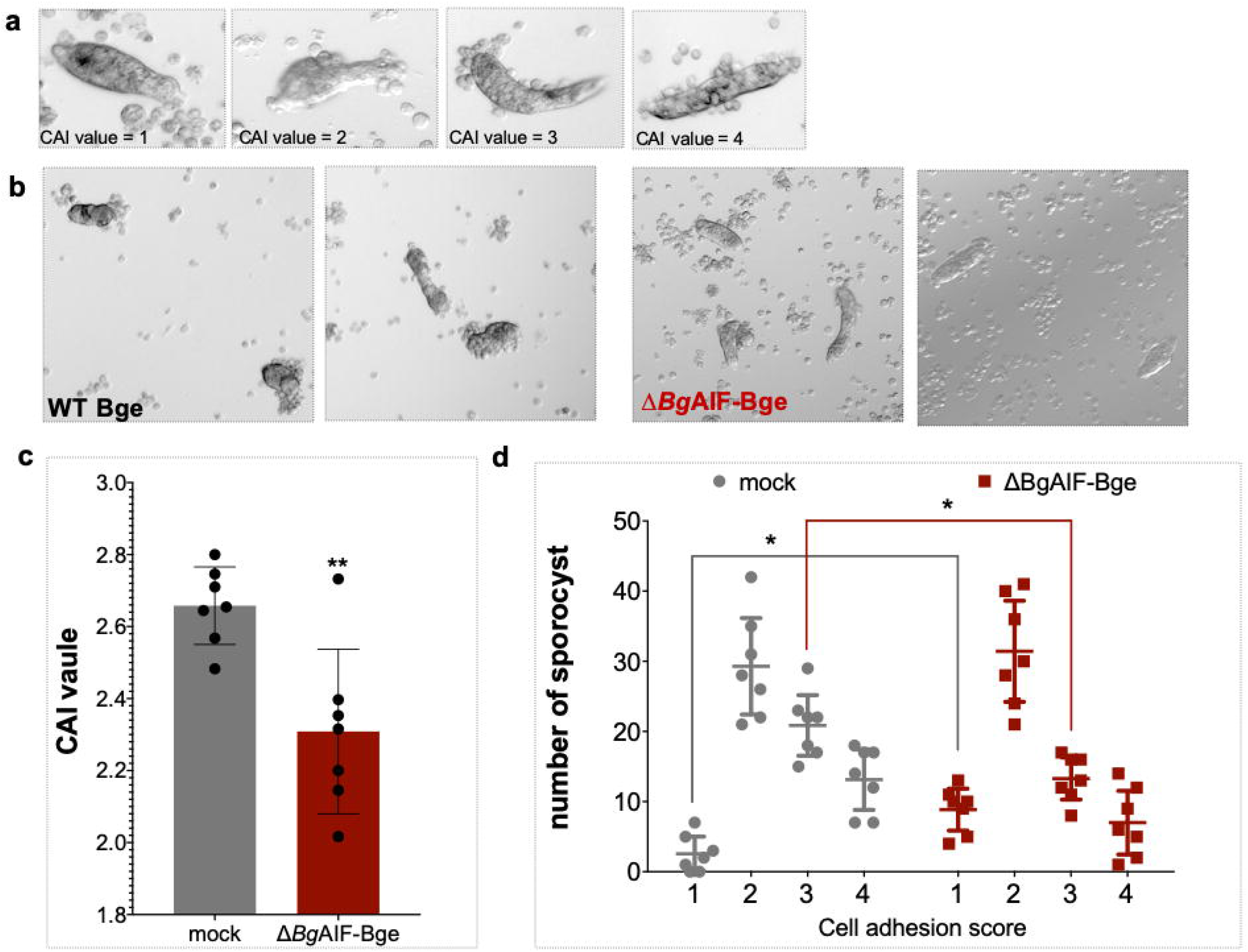
Programmed knockout of *Bg*AIF in Bge cells caused reduced adherence to primary sporocysts. **a** Representative micrographs of primary sporocysts co-cultured with Bge cells in our laboratory to profile the semi-quantitative scoring of the cell adhesion index (CAI); CAI value = 1; no cells adhering to the surface of the sporocyst; value = 2; ≤10 cells adhering to the sporocyst; value = 3; > 10 cells < half of the sporocyst surface covered by cells or clumps of cells; value = 4; > half the sporocyst surface covered by Bge cells. **b** Representative micrographs indicate the reduced levels of Δ*Bg*AIF-Bge cells adherence (right panel) in comparison to control, mock-transfected Bge cells (left panel) to the co-cultured sporocysts. **c** Bar chart to present the CAI values from control (mock-transfected) Δ*Bg*AIF-Bge cells during co-culture with primary sporocysts at a co-culture ratio of one sporocyst to 100 Bge cells; CAI value = 2.66±0.10, mean ±SD (476 sporocysts in total scored) for the mock-transfected Bge and 2.31±0.23 for the Δ*Bg*AIF-Bge cells (424 sporocysts in total scored); *P* = 0.0033, unpaired Student’s *t* test; n =7 biological replicates. **d**. The cell adhesion score ranging from 1-4 of individual sporocyst. There were 2.5±SE 1.58, 29.28±4.06, 20.85±2.15 and 13.14±2.55 for cell adhesion scores 1, 2, 3 and 4 from mock, respectively. While in Δ*Bg*AIF-Bge cell shown 8.86±1.57, 31.42±4.07, 13.28±2.51 and 13.13±2.55 of cell adhesion scores 1, 2, 3 and 4, respectively. The higher amount of sporocyst from *Bg*AIF edited-mixed cells population shown statistic significantly lower than control cell by *t-*test (t ratio = 3.98, P = 0.001). Also the lower amount of sporocysts scored as ‘3’ in Δ*Bg*AIF-Bge cells comparing with control cell (t ratio = 2.40, P = 0.004).

## Results

### Cas9 nuclease transcribed in transfected Bge cells

Total RNA was extracted from non-transfected cell (wild type; WT), mock control and pCas-*Bg*AIFx4-transfected Bge cells to assess the expression of Cas9 (Fig. 1b). The cDNAs from either controls or pCas-*Bg*AIFx4-transfected Bge cells were employed as the template in PCRs using two primer pairs, one specific for Cas9 and the other for *Bg*Actin, the actin gene of *B. glabrata* that served as the reference gene (Fig. 1b, c). Transcripts encoding Cas9 in transient pCas-*Bg*AIFx4 transfected-Bge cells were detected at 24 hrs after transfection and expression was maintained for the nine days of the assay. The specific amplicon of Cas9 mRNA (231 bp) was observed from pCas-BgAIFx4 transfected cells, but was absent from the non-transfected cells (Fig. 1c). Our findings supported previous findings that revealed s CMV promoter driven luciferase in Bge cells (60). Expression of the control reference *Bg*Actin was observed at 214 bp amplicon in both controls and experimental samples (Fig. 1c).

### Programmed mutation of *Bg*AIF confirmed functional CRISPR/Cas9 activity in Bge cells

Genomic DNAs from wild type Bge, medium-transfected (mock) and pCas-*Bg*AIFx4-transfected cells were used as the template for PCRs with the primer pair, AIF-F and AIF-R, flanking the programmed Cas9 cleavage site on *Bg*AIF, exon 4 (Fig. 1a, green arrows; amplicon size, ∼200 nt). The red arrow indicates the predicted site of the Cas9-catalyzed double-strand break (DSB) within the *Bg*AIF locus (Fig 1a). The nucleotide sequence of the amplicons was determined by Sanger direct sequencing using the same primers. Both forward and reverse Sanger direct sequencing reads from the same amplicon were estimated for insertion-deletion (INDELs) by the ICE and the TIDE algorithms (55, 56). The reads from the Bge cells transfected with the pCas9-*Bg*AIFx4 contained INDELs at or around the programmed CRISPR/Cas9 cleavage site. The percentage of reads that included INDELs ranged from 8.9% to 17.1%, in the 12 biological replicates that were carried out (Fig. 2a, b). Notably, the mutation profile in the vicinity of the predicted DSB in *Bg*AIF was similar among these 12 replicates, which were undertaken independently. Commonly observed+ INDELs at the DSBs site as revealed by the ICE analysis included deletions of 8 to 30 bp and insertions of 1 or 2 bp (Fig. 2a). These mutations were predicted to result in frameshift mutations, the consequent loss of the open reading frame, and hence and permanent knockout of *Bg*AIF in the gene-edited Bge cell. The profile of the frequency of mutations observed in each biological replicate was used to plot the curve (Prism 8 software) presented in figure 2b. These findings demonstrated that programmed genome editing using CRISPR/Cas9 was active in Bge cells, and that the non-homologous end-joining (NHEJ) pathway (61) was active in *B. glabrata* for the repair of programmed double-stranded breaks, leading to targeted gene knockout.

### Programmed mutation interrupted expression of *Bg*AIF

The aims of the study included the investigation of the activity or not CRISPR/Cas9 gene editing in the Bge cell line and addressing the hypothesis that AIF functions in the activation of a macrophage like phenotype by the Bge cell. Accordingly, Bge cells were transfected with pCas9-*Bg*AIFx4 plasmid DNA. The experimental approach did not include drug resistance and/or reporter gene markers in order to enrich for transfected Bge cells. However, even without enrichment for transfected cells, there was a highly statistically significant reduction in levels of *Bg*AIF transcripts in the transfected Bge cell population. Expression of *Bg*AIF transcripts were assessed using RNAs from the cells at nine days post transfection. Comparison of the experimental and control groups revealed significantly reduced levels of the *Bg*AIF in the pCas-*Bg*AIFx4-transfected cells, mean 49.55±20.22%, range 28.1 to 86.3% (n =12) compared to the wild type Bge (normalized sample, 100% expression), mock control cells (unpaired *t*-test and *F* test to compare variances; F, DFn, Dfd=294.7, 11, 11, *P* < 0.0001) (Fig. 1b). An inverse correlation between the percentage of INDELs and reduction in transcript levels was not apparent (not shown).

### Programmed knockout of *Bg*AIF interfered with adherence of Bge cells to schistosome sporocysts

Single cell suspensions of Bge cells in the mock-treated and Δ*Bg*AIF groups were co-cultured for 24 hrs in siliconized tubes with primary *S. mansoni* sporocysts. At that point, the numbers of cells that had adhered to each sporocyst were scored. This was accomplished by examination of at least five discrete sites of the well of the 24-well plate with ≥50 sporocysts of each group. The cell adhesion index (CAI) were scored from 1 to 4, with a score of 1 indicating few or no adherent cells and a score of 4 indicating that cells or clumps of cells covered more than half the tegumental surface of the sporocyst, as defined in earlier reports (26) (Fig. 3a). Cells from mock-transfected control group mostly adhered in clumps or singly to the surface of the parasite (representative images in the upper panels of Fig. 3B), with CAI values that ranged from 2 to 4. By contrast, fewer cells adhered to the surface of the sporocysts in the Δ*Bg*AIF-Bge group (representative images, lower panels in Fig. 3B), with CAI values ranging from 2 to 3. Only ∼20% of the Δ*Bg*AIF-Bge cells adhered to the surface of the sporocysts and most of the cells retained remained spread singly on the surface of the well of tissue culture plate (Fig. 3b). More specifically, the mean CAI values ascertained from the seven biological replicates (≥50 parasites in each replicate (≥400 parasites scored), mean 2.66±0.10, range, 2.53 to 2.78 in the mock-treated, transfection control group was significantly higher than the Δ*Bg*AIF group, mean 2.25±0.22, range, 2.08 to 2.55 (Fig 3c) (unpaired *t-*test: t=3.661 df = 12, *P* = 0.0033). More specifically, the CAI category-specific CAI values for mock treated cells averaged from the cell adherence to single sporocysts, were 2.5±1.58 SE, 29.28±4.06, 20.85±2.15, and 13.14±2.55 for categories 1, 2, 3 and 4, respectively. For the Δ*Bg*AIF-Bge cells, the CAI values were 8.86±1.57, 31.42±4.07, 13.28±2.51, and 13.13±2.55 for categories 1, 2, 3 and 4, respectively.

Although we observed CAI scores of 1 to 4 in both mock control cells and Δ*Bg*AIF cells, nonetheless there were statistically significant higher numbers of sporocysts with the lowest adherence, scored as ‘1’, in the Δ*Bg*AIF group compared with the mock control group as confirmed using a multiple *t*-test (*t* ratio = 3.98, df = 12, *P* = 0.001). By contrast, there were significantly higher numbers of sporocysts scored as ‘3’ in the control compared with the Δ*Bg*AIF group (*t* ratio = 3.52032, df = 12, *P* = 0.004) (Fig. 3d). Last, morphological changes were not apparent between the Δ*Bg*AIF and the control group Bge cells.

## Discussion

This report describes a novel use of programmed genome editing by the CRISPR/Cas9 approach in the embryonic cell line from the gastropod snail, *B. glabrata*, an intermediate host snail of the human blood fluke, *S. mansoni*. The Bge cell line is an informative tool in investigation of snail-schistosome, host-parasite interactions. A key attribute of the Bge cell is its hemocyte-like phenotype, given the central role of the snail hemocyte in innate and cellular immunity.

However, even though Bge cells adhere to the schistosome, the parasite is not killed by these cells *in vitro*. The allograft inflammatory factor 1 (AIF) is a conserved calcium-binding protein typically expressed in phagocytic and granular leukocytes and is a marker of macrophage activation (38, 41, 45, 62-65). An orthologue, termed *Bg*AIF, is highly expressed in isolates of *B. glabrata* that are resistant to infection with *S. mansoni* and this gene may be linked to hemocyte activation (8, 9). Here, we targeted the *AIF* gene of *B. glabrata* embryonic cell line using programmed gene knockout to further interrogate its role in the intermediate host-schistosome interaction. We constructed a plasmid vector encoding the CRISPR/Cas9 nuclease and a guide RNA targeting exon 4 of *Bg*AIF gene and the Cas9 nuclease from *Streptococcus pyogenes*. Bge cells were transfected with the gene-editing construct by square wave electroporation. Transcript levels of *Bg*AIF were significantly reduced by up to 71.9% following transformation. In parallel, sequence reads of amplicons spanning the locus targeted for programmed gene knock-out revealed on-target mutation on the *Bg*AIF gene, that had been repaired by non-homologous end joining leading to gene-inactivating insertions and deletions. In addition, the adherence of gene-edited Bge cells to sporocysts was significantly impeded in comparison to control cells, as ascertained using a semi-quantitative, cell adherence index. In our study, the %INDELs (8.9-17.1%) resulting from NHEJ after CRISPR/Cas9 gene editing on *Bg*AIF exon 4 locus did not correlate with its transcript reduction (∼50%) in all experimental samples. Nonetheless, alternative mechanisms could be used as the fusion of suppressors with a ‘dead’ Cas9 which enables gene regulation and increase the level of repression of the target gene (66).

The *B. glabrata* IDS comprises hemocytes and soluble proteins found in the hemolymph, among them the *Bg*AIF (67-69). The response of resistant mollusks is given by the adherence and encapsulation of sporocysts by hemocytes, leading to the parasite destruction (70). The AIF-1 was demonstrated to be a pro-inflammatory cytokine that regulates immune-related genes of the oyster, *Crassostrea ariakensis* (36). An orthologue in the leech *Hirudo medicinalis* promotes macrophage-like migration by a chemotactic activity, in addition to being involved in the innate immune responses as also seen in other species (41). The adherence of the mixed populations of *Bg*AIF gene-edited/non edited-Bge cells to sporocysts was significantly impeded in comparison to control cells, as ascertained using a semi-quantitative cell adherence index. These cells, albeit in a low percentage, are less responsive to the *S. mansoni* parasite. These data suggested that, in the presence of *S. mansoni*, the Bge cells need to secrete *Bg*AIF for activating the recruitment of more adherent Bge cells. Thus, the *Bg*AIF protein appears to play a role in cell recognition, migration, and/or adhesion, and to participate in the early immune response to the parasite. The AIF gene is conserved broadly among protostomes and deuterostomes, including vertebrates, and also in prebilaterian including sponges, where it likely performs similar functional roles in macrophage activation and migration (71). In humans, the *Hm*AIF1 is an NF-κB pathway regulator, a pathway that comprises a family of evolutionarily conserved proteins, important to the immune system by participating in the expression of other proteins related to the immune system (72, 73). Although more studies will be required to decipher the regulation of these pathways in *B. glabrata*, after the pathogen invasion, the *Bg*AIF possibly acts throughout the activation of the NF-κB pathway, leading to the recruitment of hemocytes and consequent pathogen elimination (72, 74).

These findings confirmed the tractability of transfection of Bge cells by electroporation with the genome-editing construct, pCas-*Bg*AIFx4, and that the CMV promoter drove transcription of Cas9 in this snail species. Whereas transformation by plasmid DNA of Bge cells by square wave electroporation appears to be novel, Bge cells have been transformed using DNA complexed with cationic lipid-based transfection reagents and with polyethyleneimine (23) Nevertheless, our study has some limitations. Thus far we have yet to enrich the transfected cells from wild type cells. Future studies using a drug selectable marker can be designed to address this issue. Other approaches to deliver the CRISPR/Cas gene-editing cargo can be tried including repeated inoculation with ribonuclear protein complexes (75), titration of the transfection chemicals (76), titration of electroporation parameters (77), and/or transduction by lentiviral virions encoding the gRNA and *S. pyogenes* Cas9 nuclease as we have demonstrated with eggs *of S. mansoni* (53, 54). Moreover, CIRCLE-Seq and like approaches can be employed to investigate the off-target mutations (78).

### Conclusions

Here we demonstrated CRISPR/Cas-based gene editing in a cell line from a medically important taxon of freshwater gastropods that are vectors for the transmission of schistosomiasis. We showed the functional role of a *B. glabrata* allograft inflammatory factor in the recognition/attachment of *S. mansoni* sporocysts *in vitro*. The demonstration of the activity of CRISPR/Cas9 gene editing in Bge cells suggests that genome editing in the germline and somatic tissues of intact *B. glabrata* snails will also be functional. Whereas improvements can be anticipated in these approaches, an obvious next step will be to gene edit the intact snail *B. glabrata*. Transfection of germline cells within the snail using microinjection can be considered (79). These findings, together with the first application of the CRISPR/Cas technique in the genetic edition of *Lymnaea stagnalis* mollusk (48) are a step-change since they can favor the creation of a genetically modified *Biomphalaria* line to study the biology and physiology of the snail as well the schistosome-intermediate host relationship. Functional genomics using CRISPR/Cas-based genome editing in schistosomes and other trematodes responsible for major neglected tropical diseases has been reported (53, 54). The establishment of a functional genomic protocols involving programmed gene editing to address fundamental questions in this host-parasite relationship using genetically modified snails and schistosomes now seems to be feasible.

## Abbreviations

AIF: allograft inflammatory factor 1
Bge: *Biomphalaria glabrata* embryonic cell line
Cas9: CRISPR associated protein 9
RNA: ribonucleic acid
PCR: polymerase chain reaction
CAI: cell adherence index
CRISPR: Clustered Regularly-Interspaced Short Palindromic Repeats
gRNA: guide RNA
CMV: cytomegalovirus
DBS: double strand break
SRC: Schistosomiasis Resource Center
BRI: Biomedical Research Institute
RT-PCR: reverse transcription PCR
DNA: deoxyribonucleic acid
cDNA: complementary deoxyribonucleic acid
IDS: internal defense system
EDTA: ethylenediaminetetraacetic acid
INDEL: insertion–deletion mutation
TIDE: Tracking of Indels by Decomposition
ICE: Inference of CRISPR Edits analysis
DNase I: deoxyribonuclease I
CBSS: Chernin’s balanced salt solution
WT: wild type
NHEJ: non-homologous end-joining
NF-κB: nuclear factor kappa-B

## Acknowledgments

We thank Dr. Mathilde Knight for informative discussions on the snail-schistosome immunobiology and Dr. Margaret Mentink-Kane for support with the Bge cell line. Bge cells were provided by the NIAID Schistosomiasis Resource Center of the Biomedical Research Institute, Rockville, Maryland through NIH-NIAID contract HHSN272201000005I for distribution through BEI Resources.

## Ethics approval and consent to participate

The protocols and procedures to maintain *S. mansoni* life cycle in mice performed at the NIAID Schistosomiasis Resource Center of the Biomedical Research Institute, Rockville were approved by the Institutional Animal Care and Use Committee (IACUC) (protocol number 18-04) and followed by United States Animal Welfare Act and George Washington University IACUC policies (assurance number A3205-01)

## Consent for publication

Not applicable

## Availability of data and materials

Data supporting the conclusions of this article are included within the article. The raw datasets used and analyzed during the present study are available from the corresponding authors upon reasonable request.

## Authors’ contributions

WI, MM and PJB designed the study. FSC and WI wrote the protocol. MMM, VMH and PJB reviewed the protocol. WI served as study team leader and director. VHM and AM obtained ethical approval for vertebrate animal use. FSC, RR, AM and SEK conducted the gene editing and cell culture experiments. FSC, RR and WI performed gene mutation analysis. FSC, OSC and RCL performed CAI analysis. WI, MMM and PJB wrote the manuscript. All authors read and approved the final manuscript.

